# Social intelligence model with multiple internal models

**DOI:** 10.1101/285353

**Authors:** Takuya Isomura, Thomas Parr, Karl Friston

## Abstract

To exhibit social intelligence, animals have to recognize who they are communicating with. One way to make this inference is to select among multiple internal generative models of each conspecific. This induces an interesting problem: when receiving sensory input generated by a particular conspecific, how does an animal know which internal model to update? We consider a theoretical and neurobiologically plausible solution that enables inference and learning under multiple generative models by integrating active inference and (online) Bayesian model selection. This scheme fits sensory inputs under each generative model. Model parameters are then updated in proportion to the probability it could have generated the current input (i.e., model evidence). We show that a synthetic bird who employs the proposed scheme successfully learns and distinguishes (real zebra finch) birdsongs generated by several different birds. These results highlight the utility of having multiple internal models to make inferences in complicated social environments.

## INTRODUCTION

One of the most impressive abilities of biological creatures is their capacity to adapt their behaviour to different contexts and environments (i.e., cognitive flexibility) [1,2] through unsupervised learning [3,4]. For example, people can learn to call on various responses depending on the situation, independently move the right and left hands when playing an instrument, and speak several different languages (multilingual). Such a multi-tasking ability is particularly crucial in an environment that includes several different people, who each demand subtly different forms of interaction [5,6].

Experimental studies using primates have shown that the volumes of certain brain structures (e.g., hippocampus) are correlated with the performance of cognitive and social tasks [7,8], and that the ability to infer another’s intentions increases with brain volume [9]. These findings from comparative neuroanatomy suggest that as the brain becomes larger, it can entertain more hypotheses, or internal models, about how its sensations were caused. This speaks to a putative strategy for making inferences about several different conspecifics with a number of internal models, each of which is associated with one member of their community.

This ability of biological creatures contrasts with current notions of artificial general intelligence. The development of a synthetic system –– as flexible as the biological brain –– remains an outstanding challenge [10,11]. Recent progress in this area is the introduction of a mechanism that protects parameters that have been useful in a previous context [12,13]. This enables the learning of a new task using parameters that have not been previously used, while retaining those that have been useful in the past. We draw upon this idea to try to understand how the brain might maintain distinct generative models for different contexts. To do this, we focus upon a social task (communication through birdsong) in which the conversational partner may change. This induces the dual task of inferring the identity of a conspecific, and learning about them at the same time. Crucially, this learning should be specific to each conversational partner.

To address this problem, we appeal to generalized Bayesian filtering, a corollary of the free-energy principle [14,15]. We illustrate the proposed scheme using artificial birdsongs and natural zebra finch songs. We present a synthetic (student) bird with a song generated by one of several (teacher) conspecifics. During the exchange, the student bird performs Bayesian model selection [16] to decide which teacher generated the heard song. Having accumulated sensory evidence under all hypotheses or models, the parameters of the generative models are updated in proportion to the evidence for each competing model. We show that, over successive interactions, our student is able to learn the individual characteristics of multiple teachers –– and recognise them with increasing confidence.

### Concept of modelling

In formulating the generative model, we have to contend with a mixture of random variables in continuous time (i.e., latent states of each singing bird) and categorical variables (i.e., the identity of the bird) that constitute a perceptual categorisation problem. In short, the listening bird (i.e., student) has to make inferences in terms of beliefs over both continuous and discrete random variables in order to recognise *who* is singing and *what* they are singing. In a general setting, this would call upon mixed generative models with a mixture of continuous and discrete states –– of the sort considered in [17] and more recently [18]. A complementary way of combining categorical and continuous latent states is to work within a continuous generative model that includes switching variables that have a discrete (i.e., categorical) probability distribution, with an accompanying conjugate prior such as the Dirichlet distribution. The most common example of this would be a Gaussian mixture model: see [17] for details.

Heuristically, this means the generative model can be constructed in one of two ways. We can either select a singing bird to generate a song; leading to a deep model with a categorical latent variable at the top and a continuous model generating outcomes. Alternatively, we could generate continuous outcomes from all possible birds and then select one to constitute the actual stimulus. In the second (switching variable) case, the categorical variable plays the role of a switch; basically, switching from one possible sensory ‘channel’ to another.

In terms of model inversion and belief propagation, both generative models are isomorphic –– and lead to the same update equations via minimisation of variational free energy. However, the way in which the generative models play out, in terms of requisite message passing, can have different forms. We could use a generative model with a single bird and try to infer which bird was singing (and, implicitly, the parameters of its generative process). Alternatively, the student may entertain all possible teachers ‘in mind’ and then select the best hypothesis or explanation for the sensory input. This would correspond to the second formulation of the generative model, in which the dynamics are conditioned upon the categorical variable (i.e., songs are simulated under all possible hypotheses) and the best explanation is then selected. In this sense, the expectation about the identity of the singing bird acquires two complementary interpretations. In the first formulation, it is the posterior expectation about the bird that has been selected to generate the song. In the second interpretation, it becomes an expectation about the switching variable. This means the student (i.e., listening bird) effectively composes a Bayesian model average over all hypotheses (i.e., singing birds) entertained, in providing posterior predictions of the song.

We can appeal to both interpretations when interpreting the results below. However, the second interpretation has some interesting interpretations from a cognitive neuroscience perspective. In essence, the gating or selection of top-down predictions complements the gating or selection of ascending prediction errors usually associated with attention [19–21]. In other words, selecting (switching to) the best explanation from available hypotheses –– when predicting sensory input –– becomes a covert form of (mental) action. A famous example of such an attentional switching is visual illusion [22]. This is in the sense that descending predictions are contextualised and selected on the basis of higher order beliefs (i.e., expectations) about the most plausible hypothesis or context in play. The unique aspect of this gating rests upon the fact that there are a discrete (categorical) number of competing hypotheses, which are mutually exclusive. This is reminiscent of equivalent architectures in motor control (e.g., the MOSAIC architecture) and related mixture of experts [17,23]. In our case, a simple perceptual categorisation paradigm mandates a selection among different possible categories and enforces a form of ‘mental action’ through optimisation of an implicit switching variable.

In what follows, we present the results of perceptual learning and inference using this form of model selection or *structure learning*, predicated on an ensemble or repertoire of generative models (using synthetic birds and real bird songs). In this setting, we show that Bayesian model averaging provides a plausible account of how multiple hypotheses can be combined to predict the sensorium, while Bayesian model selection enables perceptual categorisation and selective learning (i.e., without forgetting). Crucially, all of these unsupervised processes conform to the same normative principle; namely, the minimisation of (the path integral of) a variational free energy bound on model evidence.

## RESULTS

### Multiple generative models and attentional switching

Organisms continuously infer the causes of their sensations (e.g., unconscious inference or Bayesian brain hypothesis) [24,25] and thereby predict what will happen in the immediate future (e.g., predictive coding) [26,27]. This sort of perceptual inference rests upon an internal generative model that expresses beliefs about how sensory inputs are generated. These models typically assume that sensations are generated by latent or hidden (unobservable) causes in the external world. Such causes may themselves be generated by other causes in a hierarchical manner. In the setting of continuous state-space models, hierarchical Bayesian filtering can be used to perform inference under a hierarchical generative model [14,28]. This filtering uses variational message passing to furnish approximate posterior probability (recognition) densities over the hidden states. In what follows, we describe the process generating sensory inputs. We consider that the same model structure is also used in the brain as an internal generative model; however, the brain does not know the values of hidden states and underlying model parameters [29–31]. A detailed description of the generative models used in this study is provided in Supplementary Methods S1. See also Supplementary Table S1 for glossary of expressions.

Let us consider a birdsong generative model as an example. In brief, this model is a deep generative model with two levels; both based upon chaotic dynamics in the form of Lorenz attractors (see Methods): see also [32,33] for details. Crucially, the state of a (slow) higher attractor (associated with neuronal dynamics in Area X in the songbird brain) provides a control (Rayleigh) parameter for a (fast) lower attractor. The hidden or latent states of the lower attractor (associated with the higher vocal centre for HVC) then drive fluctuations in the amplitude and frequency of birdsong: see also [34–36] for related songbird studies.

In our case, we are interested in multiple models (i.e., multiple teachers) each specified by *m*_*i*_ with *i* = 1,2, … ∈ ℳ (Fig. 1 right). Each *m*_*i*_ indicates a specific model structure including certain forms of functions and dimensions of latent variables and parameters. These models describe how sensory input (i.e., birdsong) *s* is generated by a set of latent variables *u*^(*i*)^ that include hidden states *x*^(*i*)^ and hidden causes *v*^(*i*)^. Note that bracketed superscript (*i*) indicates they belong to model *i* (or bird *i*). These variables are associated with trajectories that are specified in generalised coordinates of motion 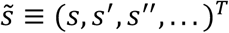, where dashes denote time derivatives. The processes that generate birdsong from these latent variables are parameterised by a set of parameters *θ*^(*i*)^. We can represent the generation of birdsong under model *i* by the following stochastic differential (Langevin) equations:

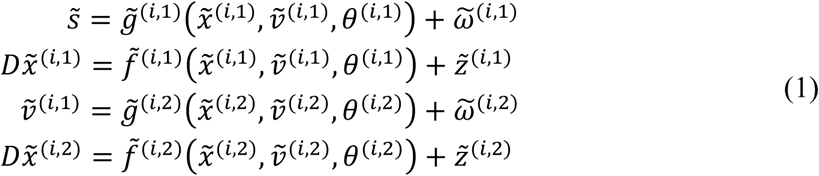

**Figure 1.**
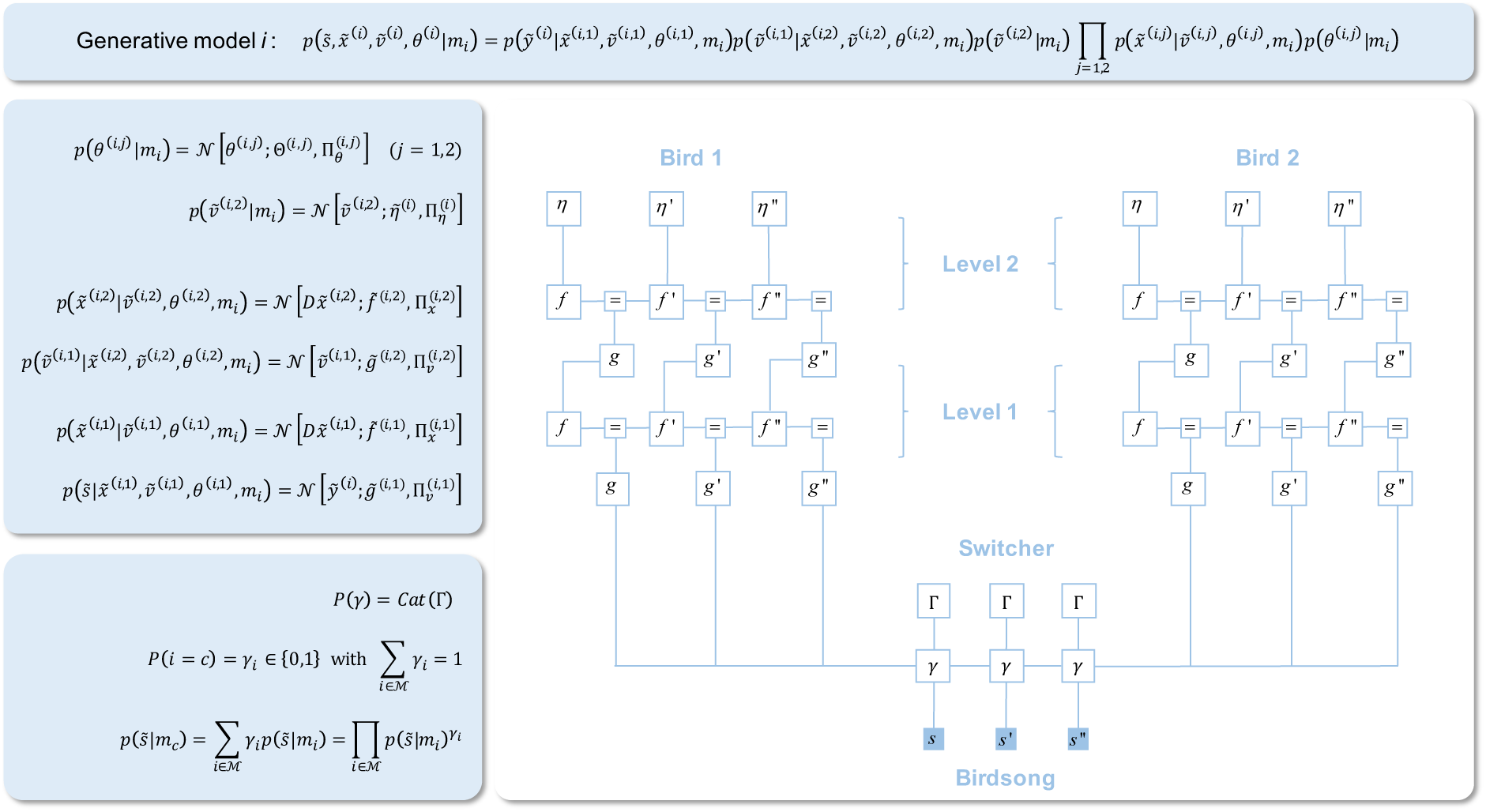
This figure illustrates how the random (stochastic) differential equations of motion (Eq. (1)) can be interpreted as a probabilistic generative model. This consists of a joint distribution (upper panel) that can be factorised into the distributions shown in the middle left panel. The large lower right panel depicts two generative models graphically, in the form of a normal (Forney) factor graph [18,76,77]. This shows that the sensory input (birdsong) may be generated by one of two teacher birds, each represented by its own hierarchical generative model. A switcher placed in the centre determines which bird generates the sensory input as described in the bottom left panel. The bottom left panel shows that the switcher state *γ*_*i*_ corresponds to the probability of model *i* being selected, where only *γ*_*c*_ takes a value of one (i.e., *m*_*c*_ is the present model), while the remaining *γ*_*i*_s with *i* ≠ *c* are zero. Importantly, regardless of the switcher state, all models generate dynamics –– e.g., 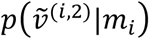: indicates the probability of the hidden (generalised motion of) cause 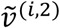, under the *i*-th model structure *m*_*i*_, while the selected model is denoted by *m*_*c*_. The task of our synthetic (student) bird is to infer which (teacher) bird generated the song (i.e., to infer *γ*_*i*_). Having done so, the parameters *θ*_*i*_ associated with bird *i* are updated in proportion to the evidence that bird *i* was, in fact, singing. See also Supplementary Methods S1 for the details.

In the above, 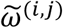 and 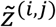 represent random fluctuations. They follow Gaussian distributions with mean zero and precision (inverse covariance) of 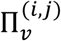 and 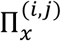, respectively. *D* is a matrix operator that implements 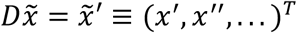. The superscript (*i,j*) indicates the *j*-th level of model *i* (i.e., latent variables at level two generate latent variables at level one that generate sensory input). The model specified by these equations can be interpreted in terms of a set of probability distributions (Fig. 1 middle left), and their product provides the *i*th generative model (Fig. 1 top).

In our task design, although every generative model is running simultaneously, only the signal generated by a specific bird is selected as the sensory input (i.e., a teacher song) that a student can actually hear –– as an analogy to social communication with several different conspecifics. This selection is controlled by a switcher (see also Fig. 1 right): suppose the currently selected model is indexed by *c*. We represent the switcher state by a set of binary variables *P*(*i* = *c*) = *γ*_*i*_ ∈ {0,1} where only *γ*_*c*_ = 1, while the remaining variables are zero, to ensure Σ_4∈ℳ_ *γ*_*i*_ = 1. Note that *γ*_*i*_ indicates the probability of model *i* being selected but it takes only either 0 or 1 by design. Using this switching process, the probability of sensory input is 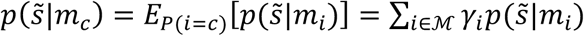, where 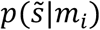 is the conditional probability of the sensory input when model *i* is selected (Fig. 1 bottom left). We suppose that the switcher *γ* is sampled from a categorical prior distribution *P*(*γ*) = *Cat*(Γ) for each epoch of singing. In sum, all models (*m*_*i*_) are generating dynamics, while only the output from *m*_*c*_ is selected as the sensory input.

### Update rules for inference, model selection, and learning

The inversion of a generative model corresponds to inferring the unknown variables and parameters that we will treat as perceptual inference and learning respectively. Formally, in variational Bayes, this rests on optimising an approximate posterior belief over unknown quantities by minimising the variational free energy (and its path integral) under each model. This comprises the following three steps as shown in the top of Fig. 2: (1) in the inference step, latent variables under all models are updated over an epoch of birdsong; (2) in the model selection step, a softmax function of variational free action, under each model, constitutes model evidence (i.e., marginal likelihood); and (3) in the learning step, the model evidence plays the role of an adaptive learning rate when updating model parameters using a descent on variational free action. This ensures only models that are likely to be generating the bird-song (sensory data) are updated, while the remaining models retain their current parameters. In what follows, we derive the associated update rules, to illustrate their general form.

**Figure 2.**
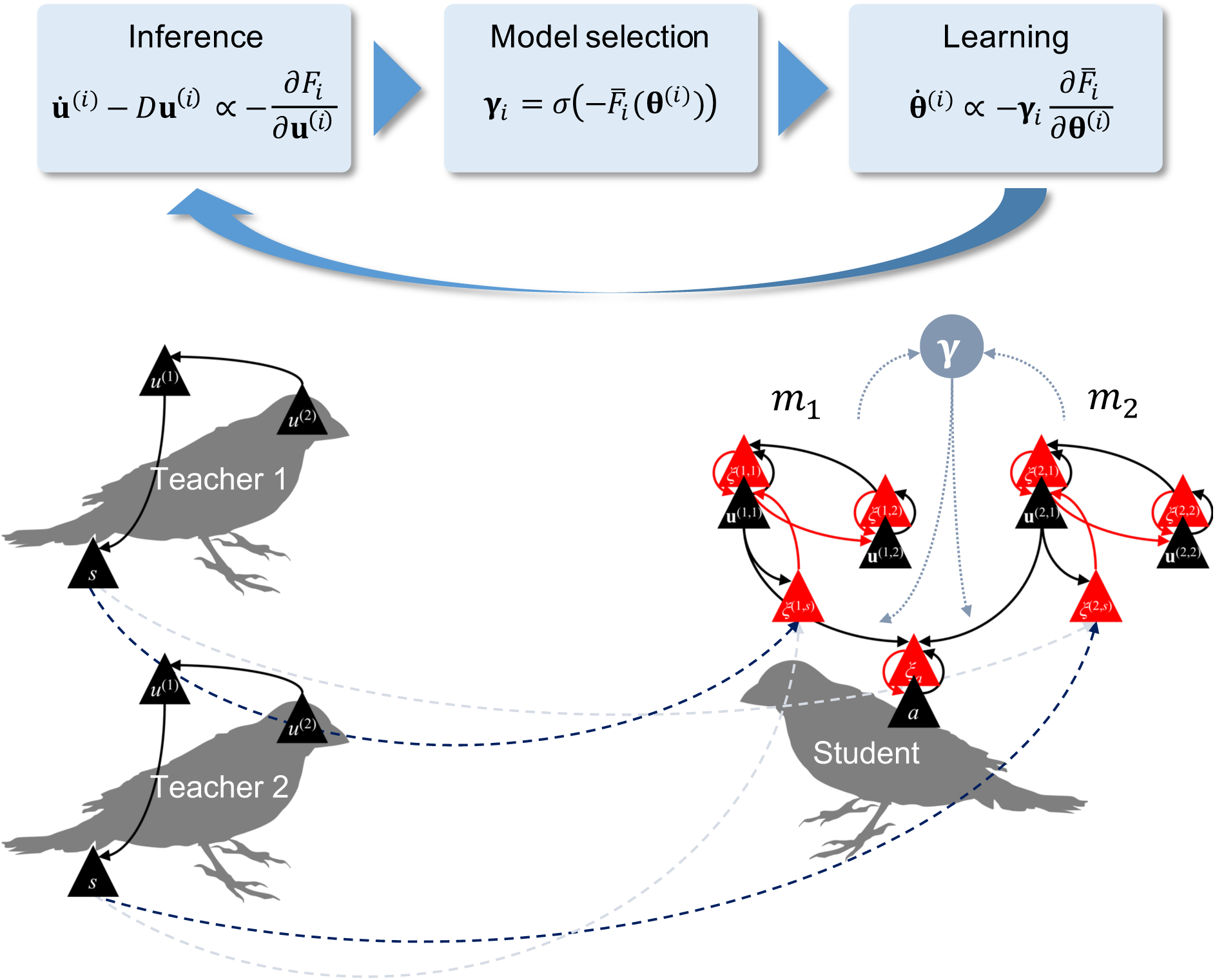
Schematics illustrating the variational update scheme (top): the models that our synthetic student (right) uses to make inferences about the songs generated by two teachers (left). A flow chart on the top summarises the inference, model selection, and learning processes that the student must implement. *σ*(·) is a softmax (normalised exponential) function that converts the conditional free actions to a model plausibility **γ**_*i*_. Our learning process is weighted by the model plausibility, ensuring that the model most likely to have generated the heard song updates its parameters during learning. See also Supplementary Methods S2 for further details.

The latent variables and parameters under model *i* are given by 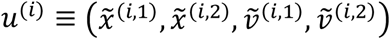 and *θ*^*i*^ ≡ (*θ*^(*i*,1)^, *θ*^(*i*,2)^), respectively. The internal energy 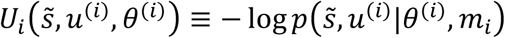 quantifies the amount of prediction error under generative model *i*, i.e., the likelihood of 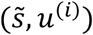 when *θ*^(*i*)^ and *m*_*i*_ are given. Using this, the conditional free energy of model *i* is given by

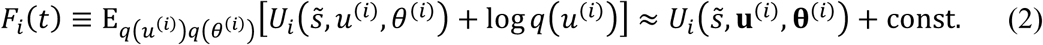

Here, *q*(*u*^(*i*)^) and *q*(*θ*^(*i*)^) are approximate posterior (i.e., recognition) densities over the latent variables and parameters of each model. The expression 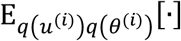 denotes the expectation over these posterior beliefs, and bold symbols **u**^(*i*)^ and **θ**^(*i*)^ denote their posterior expectations (i.e., the means of *q*(*u*^(*i*)^) and *q*(*θ*^(*i*)^)), respectively. Thus, **u**^(*i*)^ and **θ**^(*i*)^ are maximum a posteriori estimates of the latent variables and parameters under model *i*. If we ignore the second order derivative of *U*_*i*_, we can express *F*_*i*_ by simply substituting **u**^(*i*)^ and **θ**^(*i*)^ into *U*_*i*_ up to a constant term. In neurobiological process theories, **u**^(*i*)^ and **θ**^(*i*)^ are usually associated with neural activities and synaptic strengths, respectively [37].

Inference optimises the approximate posterior beliefs (expectations) about the latent variables. This can be expressed as a gradient flow in generalised coordinates of motion (noting that the solution satisfies the variational principle of least action):

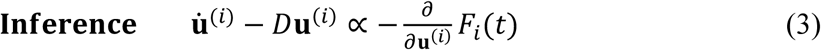

Special cases of this generalised Bayesian filtering reduce to Kalman filtering. The implicit optimisation of **u**^(*i*)^ allows for inference to take place under every model. In addition to this, our agent needs to infer which model is currently generating its sensory input. This involves minimisation of the free action (denoted by a bar) over models, given by:

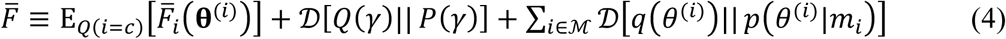

Note that the first term is the weighted sum of the path integral of the conditional free energies, where 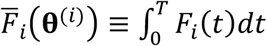 is the conditional free action of model *i*. In this expression, *Q*(*i* = *c*) = **γ**_*i*_ ∈ [0,1] with Σ_4∈ℳ_ **γ**_*i*_ = 1 denotes the posterior expectation about model *i* being selected, which is equivalent to the posterior belief about the switcher state *Q*(*γ*) = *Cat*(**γ**). The second and third terms are complexity terms relating to the switcher and parameters (expressed by Kullback-Leibler divergence [38]). When the prior distribution of the switcher state *P*(*γ*) is the same for each model (i.e., all birds are equally likely), we obtain the posterior expectation of the switcher state **γ**_*i*_ that minimises the total free action as

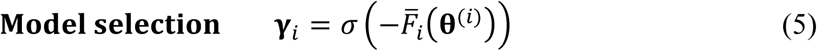

This means that the posterior expectation (i.e., evidence) that model *i* generated the song (denoted by **γ**_*i*_) can be computed by taking a softmax *σ*(·) (normalised exponential) of the conditional free actions for each model –– analogous to a *post hoc* Bayesian model selection [39] and a discrete categorical model [40]. We also refer to **γ**_*i*_ as the model *plausibility* since this quantifies how likely model *i* is to have generated the current sensory input.

Finally, learning entails updating posterior expectations about the parameters **θ**^(*i*)^ to minimise the total free action. Taking the gradient of the total free action with respect to the parameters furnishes the learning update rule. When the prior density of parameters *p*(*θ*^(*i*)^|*m*_*i*_) is flat for every model (i.e., no prior knowledge about parameters), this optimisation is given by the minimization of the conditional free action weighed by **γ**_*i*_:

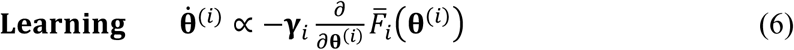

The novel aspect of this update rule is the weighting of its learning rate by the model evidence or plausibility. This means only plausible models will change their parameters, which enables the learning of several different generative models in a (soft) winner-takes-all manner. Detailed derivations of the above equations can be found in Supplementary Methods S1 and S2.

Notably, the posterior distribution of the switch *Q*(*γ*) can be considered as an attentional filter [19–21]. On this view, an attended generative model and posterior beliefs correspond to the marginal distributions over models and posteriors. When each model has the same structure and dimensions, these marginals are given by 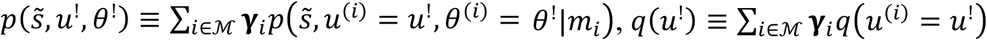, and *q*(*θ*^!^) ≡ Σ_4∈ℳ_ **γ**_*i*_*q*(*θ*^*i*^ = *θ*^!^). Thus, our model is formally analogous to a Gaussian-mixture-model version of a generalised Bayesian filter. On a more anthropomorphic note, the marginal beliefs over latent variables *u*^!^ and parameters *θ*^!^ are fictive (i.e., do not exist in the external real world). One could imagine that they underwrite some conscious inference, with several competing generative models (i.e., hypotheses) running at a subpersonal or unconscious level in the brain.

Interestingly, the above formulation can be applied to generative models that have different structures and dimensions, because there is no direct interaction between generative models – – and the switcher receives only the output from each generative model. This property may be particularly pertinent for recognising conspecifics, since conspecifics may not be best modelled using the same generative model structure.

### Demonstrations of multiple internal models using artificial and natural bird songs

A birdsong has a hierarchical structure to express complicated meanings using a finite set of notes [41]. Young songbirds are known to learn such a song by mimicking adult birds’ song [42–46]. A series of previous studies have developed a songbird model inferring the dynamics of another’s song based on a deep (two-layer) generative model [32,33]. Perceptual inference requires an internal model of how the song was generated. However, in a social situation, several birds may produce different songs generated by different brain states (or generative models). In the simulations below, we consider a case where two birds (denoted by teacher 1, 2) sang two different songs in turn, as illustrated in Fig. 2 left. A song *s* = (*s*_1_, *s*_2_)*^T^* is given by a 4-s sequence of a 2-dimensional vector, where *s*_2_ and *s*_1_ represent the mode of frequency and its power, respectively –– analogous to a physiological model of birdsong vocalizations [47]. In preliminary simulations, we confirmed that when a student with a single generative model heard their songs, it was unable to learn either teacher 1 or 2’s song (Supplementary Figures S1A–C). This is because a single generative model cannot generate two songs. Thus, the student tried to learn a spurious intermediate model of the two songs and failed to learn either.

This limitation can be overcome using a repertoire of generative models (see Methods for details). We found that a student with two generative models (*m*_1_, *m*_2_; see Fig. 2 right) can solve this unsupervised learning problem efficiently. We trained the model by providing two (alternating) teacher songs by updating an unknown parameter of both generative models. Posterior densities over parameters (i.e., synaptic strengths) were updated over learning and successfully converged to the true values used in the simulation (Fig. 3A). As a result, the student was able to make perceptual inferences about latent states generating both songs (Fig. 3B). In each session of training, both internal models were converted to produce conditional free actions. The trajectories of free action evince a process of specialisation, where each model becomes an ‘expert’ for one of two songs (Fig. 3C). At the beginning of exposure, the evidence for both models was around 0.5, which led to parameter updates in both models (Fig. 3D). Following learning, the difference in model plausibility became significantly larger –– and only the most likely model updated its parameter following the appropriate song. In Fig. 3, the hidden states of teachers were reset at the beginning of each session, which made the song sequence periodic and easy to learn. When the hidden states of teachers were not reset, the song sequence became chaotic and was more difficult to learn. However, even in this chaotic case, our model successfully learned from two distinct teachers (Supplementary Figures S2A–D).

**Figure 3.**
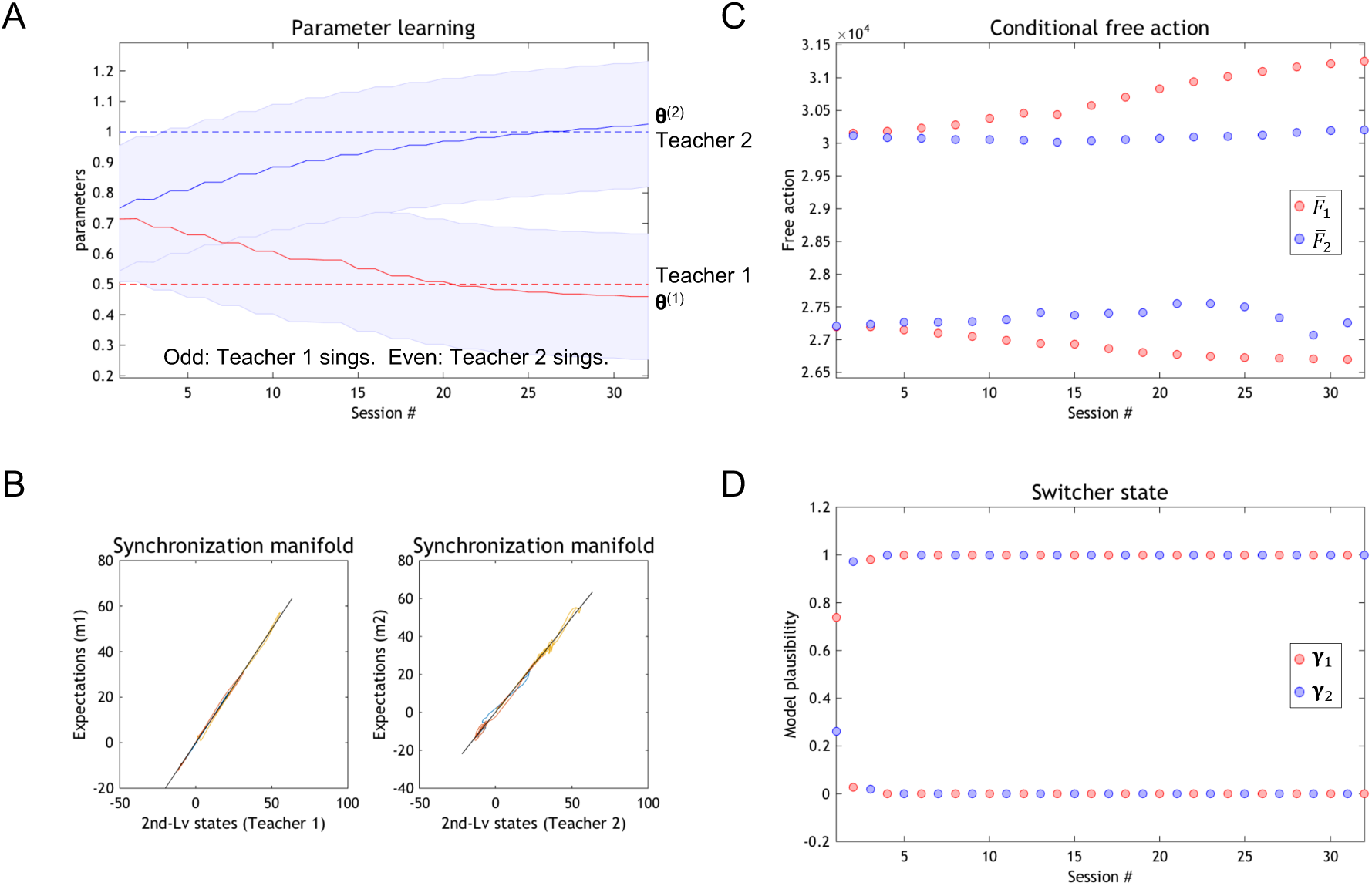
Simulation results when learning two birdsongs using multiple generative models. Teacher bird 1 generated a song in odd sessions, while teacher bird 2 sang in even sessions. At the beginning of each session, the initial latent variables of both teachers were reset to their initial values to ensure they generated quasi-periodic dynamics. The parameters of both teachers were fixed over sessions. A student bird was equipped with two generative models (*m*_1_, *m*_2_). (**A**) Trajectories of the posterior of a parameter that was optimised. The parameter for *m*_1_ and *m*_2_ (red and blue curves, respectively) were initialised from the middle point (≈ 0.75) and updated according to the variational scheme in the main text. After training, *m*_1_ and *m*_2_’s parameter approximated the true parameter value of teacher 1 (= 0.5; red dashed line) and 2 (= 1.0; blue), respectively, reflecting veridical learning. Shaded areas indicate the standard deviation of the posterior density. (**B**) Comparisons between true (teacher) hidden states and their posterior expectations inferred by the student (left: teacher 1 vs. *m*_1_, right: teacher 2 vs. *m*_2_). (**C**) Trajectories of conditional free actions for *m*_1_ (red) and *m*_2_ (blue). When teacher 1 sang (odd sessions), 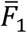 was lower than 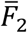, and vice versa. (**D**) Trajectories of model plausibility, used for parameter updates. MATLAB source codes for this simulation are appended as Supplementary Source Codes.

We next provided six distinct (natural zebra finch) songs to our model to see if it could learn and recognise six different teachers (Fig. 4; see also Methods for details). Before training, we tested the responses of the student to the teacher songs as a reference (Supplementary Movie S1). A student bird with six internal models inferred latent states (with a small update rate) and calculated the accompanying free energy and model evidences (Figs. 4A and 4B). After exposure, the student generated a song to predict (or imitate) the current teacher song by running the generative models in a forward or active mode. In this mode, the bird reproduces its predicted sensory input based upon a Bayesian model average of the dynamics generating a particular song. This Bayesian model average is the mixture of model specific predictions weighted by model evidence or plausibility. However, prior to learning, the student could not reproduce the teacher song because they have not learned the teacher’s parameters –– and could not categorise the teachers. During training, we randomly provided one of the six teacher songs for 60 sessions (Supplementary Movie S2). The student listened to the song and evaluated model plausibility for each of its six internal models. It then learnt (420-dimensional) unknown model parameters –– with a learning rate determined by model plausibility –– to ensure only plausible models were updated. These parameters controlled a non-linear (polynomial) mapping from latent states expressing the chaotic dynamics of the deep generative models to fluctuations in amplitude and peak frequency of the sensory input.

**Figure 4.**
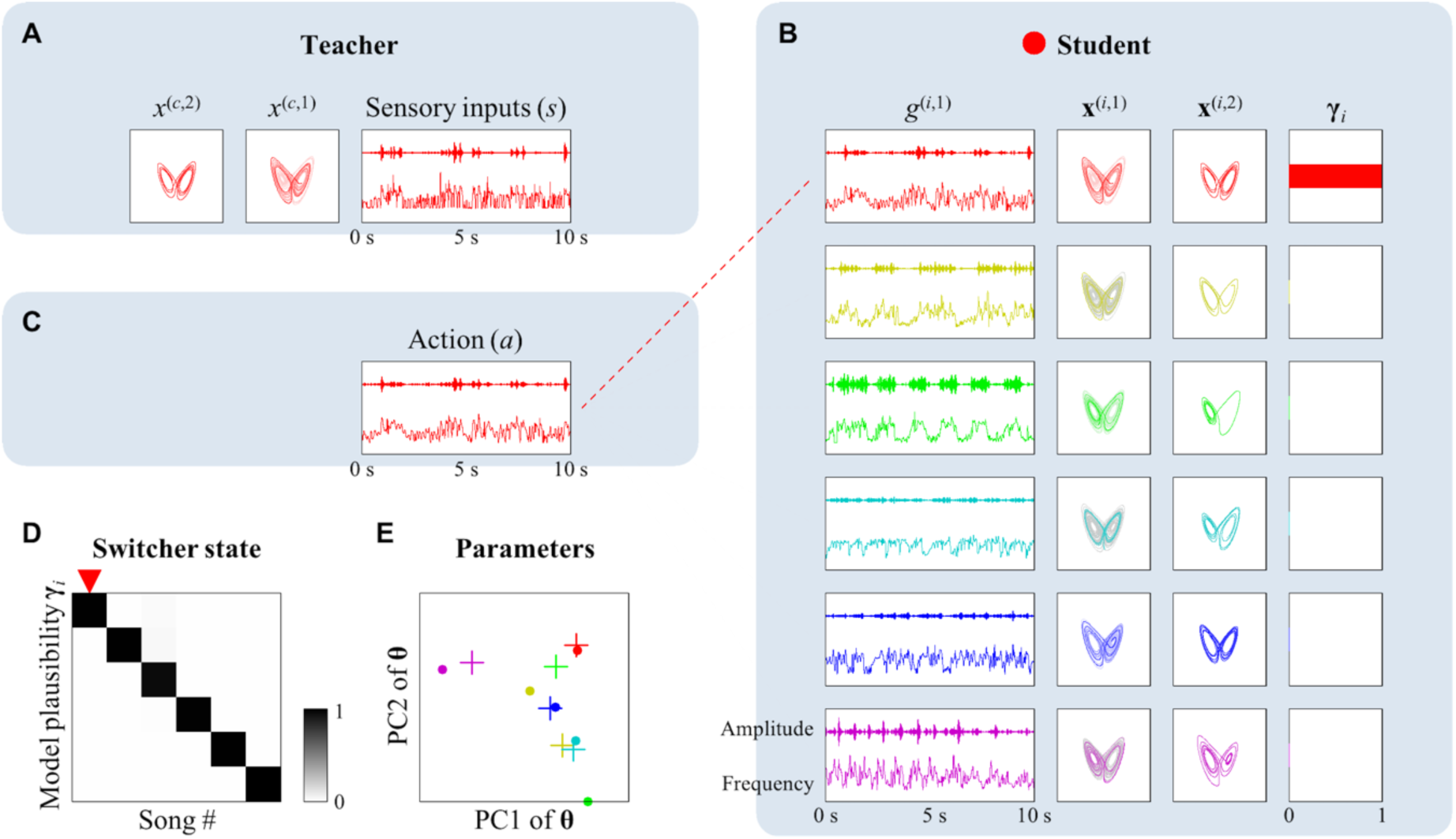
A demonstration of learning six natural zebra finch songs using the multiple generative models. The dynamics of teacher and student states before, during, and after training are provided as Supplementary Movies S1, S2, and S3, respectively. This figure shows a snap-shot of a movie after training. (**A**) A teacher song (right) and underlying dynamics of hidden states and causes (left and middle; they were estimated from the sensory data). Six real zebra finch songs were processed and used as teacher songs (illustrated in six colours). A song is given by a 10-s sequence of *s*. The currently selected song is indexed by *c*. (**B**) Six internal generative models in a student listening to a teacher, making inferences about latent states (centre), model plausibility (right), and the predicted trajectories of sensory input *g*^(*i*,1)^ (left). These constitute the expected song sequences (output) under each model. The colour of trajectories indicates the song for which each model is specialized. (**C**) To evaluate prediction capability, a student generates song (action) in the absence of sensory input. Action is given by the average of predictions of the six models weighted by model plausibility; i.e., the Bayesian model average. (**D**) Posterior expectations of the switcher state (or model plausibility) were updated in each model selection step (see Eq. (5)). They were initialised from a uniform distribution and converged to a definitive identity matrix; suggesting that each model became specialised for a specific teacher song. (**E**) Posterior expectations of the parameters of six internal models (circles) and optimal parameters for six teacher songs (plus marks) plotted in a subspace of the first and second principal components (PC1, PC2) of parameter space. During learning, only internal models with high model plausibility enjoy parameter updates (see Eq. (6)). The initial parameter values were adjusted in the absence of model selection to ensure their initial values generated an averaged song (see Methods for details).

We found that the student’s internal models became progressively specialised for one of six teacher songs (Supplementary Movie S3). After learning, only the most plausible model (with veridical parameters) contributed to the Bayesian model average, so that the student could reproduce the teacher songs in a remarkably accurate way (Fig. 4C). These results are particularly pleasing because they also suggest that real (zebra finch) songbirds generate songs (in their HVC and area X) using chaotic dynamics with the form we have assumed for previous simulations. Indeed, to compare inferred and true hidden states (and parameters), the real zebra finch songs were learned separately and regenerated under the appropriate model to provide stimuli. Learning success was further confirmed by a specialisation of each model for a specific teacher (Fig. 4D) and a convergence of posterior parameter expectations, under each model, to the teacher specific values (Fig. 4E). These results suggest that the proposed scheme works robustly, even with natural data and a large number of songs. These results highlight the potential utility of multiple generative models in learning (and inverting) highly context sensitive models of the social world.

## DISCUSSION

The brain may use multiple generative models –– and select the most plausible explanation for any given context. This strategy can be used to learn and recognise particular conspecifics in a communication all social setting. A key question here is how the brain separately establishes distinct generative models before it recognises which model is fit for purpose –– and *vice versa*. In this study, we introduce a novel learning scheme for updating the parameters of the multiple internal models that are themselves being used to filter continuous data. First, several alternative generative models run in parallel to explain the sensory input in terms of inferred latent variables; this enables the conditional free action and associated model plausibility to be evaluated; finally, the parameters of each model are updated with a learning rate that is proportional to the model plausibility. This ensures only models with high model plausibility or evidence are informed by sensory experience. The proposed scheme allows an agent to establish and maintain several different generative models (or hypotheses) and to perform an adaptive online Bayesian model selection (i.e., switching) of generative models depending on the provided input. Crucially, there is no danger of forgetting previously learned parameters and structures, because model selection and implicit structure learning is context sensitive –– and this context is inferred explicitly. Furthermore, a repertoire of generative models may be able to facilitate an adaptation speed to a new environment. In the subsequent work, we will address this issue.

Neurobiologically, our learning update rule might be implemented by associative (Hebbian) plasticity modulated by a third factor, a concept that has recently received attention [48– 50]. While Hebbian plasticity occurs depending on the spike timings of pre- and post-synaptic neurons [51–54], recent studies have reported that various neuromodulators [55–60], GABAergic inputs [61,62], and glial factors [63] can modulate Hebbian plasticity in various ways. Our learning update rule consists of the product of the (conditional) free action gradient providing a Hebbian-like term (see [14] for details) and the posterior belief of the switcher state, in which the latter might be implemented by such third neurobiological factors.

Previous studies have modelled communication between agents [64,65] in analogy with the mirror neuron system [66,67]. These simulations involve two birds who infer each other, converging onto the same internal state, and generating the same song. This has been used as a model of hermeneutics, cast in terms of generalised synchrony (or synchronisation of chaos). Heuristically, both birds come to sing from the same ‘hymn sheet’ and thereby come to ‘know each other’ through knowing themselves. Such a synchronous exchange minimises the joint free energy of both birds –– because both birds become mutually predictable. This would be related to an experimental observation that a birdsong can propagate emotional information to another bird and make influence on its behaviour [68]. This set can be generalised, when more than two birds are singing the same song. However, when several conspecifics generate different songs (or speak different languages), an agent with a single generative model is no longer fit for purpose. We addressed this limitation by equipping synthetic birds with alternative attractors or hypotheses. Interestingly, it has been reported that birdsongs have such a learning flexibility [69]. In our case, generative models are explicitly decomposed and their learning rates are tuned by an attentional switch, allowing an agent to optimise each model for each context –– the existence of neurons exhibiting such a teacher specific activity has also been reported in the songbird higher-level auditory cortex [45]. In other words, inference about a particular correspondent’s ‘state of mind’ can be modelled by synchronous dynamics during conversation, where one of listener’s internal models converges to an attractor representing the speaker’s. On this view, empathic capacity may be quantified by how many attractors the agent can deploy –– and how well it can optimise each attractor.

Moreover, we applied the scheme to infer several different natural zebra finch songs. To deal with natural birdsongs, we used a large number of model parameters and a small update rate for inference. This is conceptually analogous to reservoir networks [70–72], where an input sequence is inferred using a linear mapping from the outputs of a randomly connected chaotic neural network. In our case, we represented a birdsong using a nonlinear mapping from the dynamics of two Lorenz attractors with distinct time constants. Since each internal model generates a characteristic attractor, significant differences between model plausibility can emerge. This makes learning with multiple internal models more efficient.

Using multiple generative models, one can develop a computational model that infers another’s mind while distinguishing itself from another. For example, the ‘Sally-Anne test’ is a diagnostic paradigm designed to test for theory of mind in autism. It asks whether one can distinguish one’s own beliefs from another’s [73–75]. This form of higher-order inference can be reproduced using multiple generative models (Fig. 5). For example, a teacher is modelled as a simple birdsong generator, while student 1 has a single internal generative model to infer and imitate the teacher’s song. Student bird 2 hears two different birdsongs generated by a teacher and student 1, and establishes two generative models (Fig. 5A). One model is used to infer the teacher’s song, which expresses student 2’s belief about the environment (i.e., my mind), while the other model infers student 1’s song, expressing how student 1 perceives the world (i.e., another’s mind). The importance of having two internal models is disclosed when the teacher changes its song in the absence of student 1 (Fig. 5B). In this case, student 1 does not update its knowledge about the teacher’s song. Student 2 updates model parameters only in a model representing the teacher’s song, while maintaining the other model representing student 1. In subsequent work, we will show how the ability to distinguish between how an agent perceives the world and how another thinks underwrites theory of mind and perspective taking.

**Figure 5.**
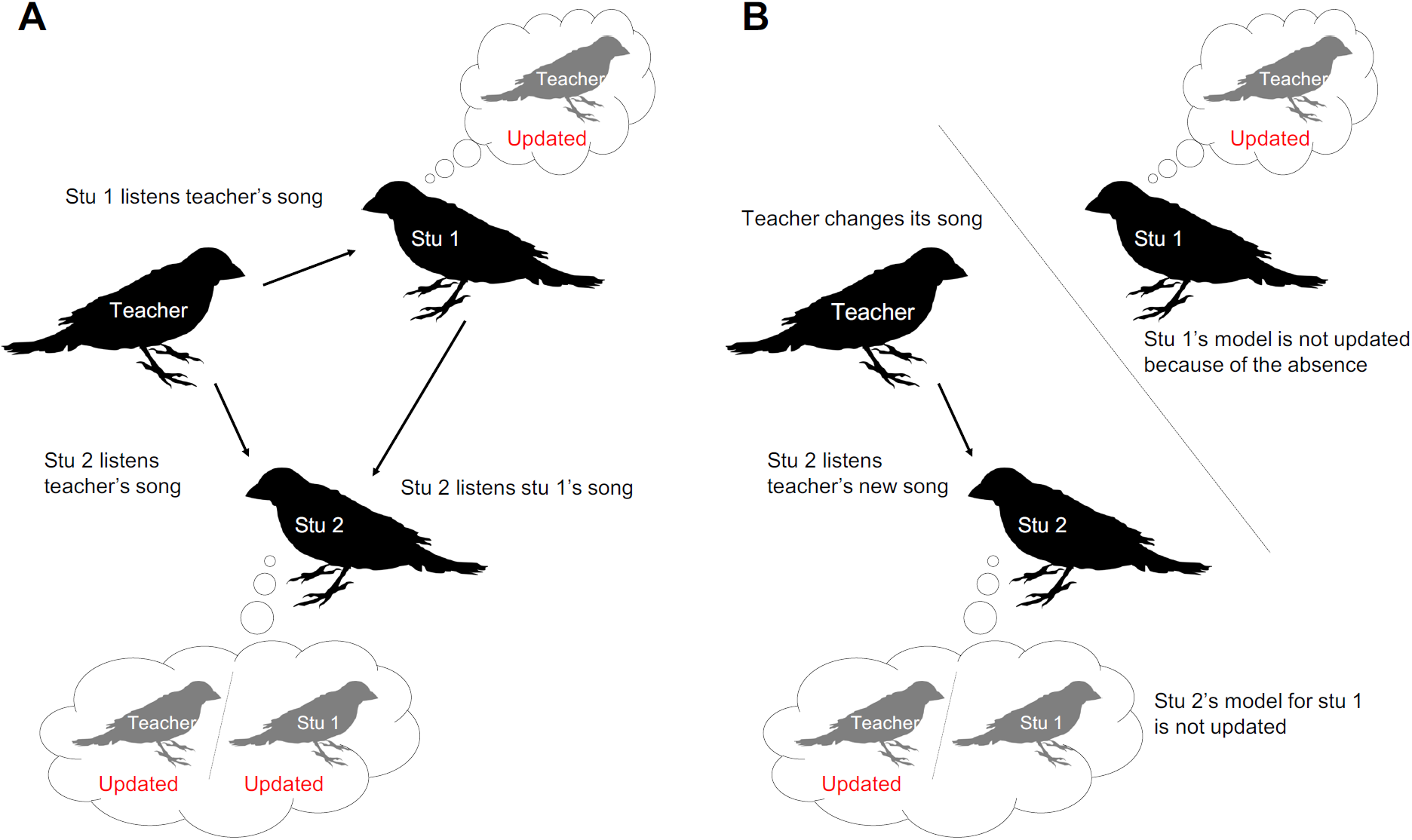
An illustration of a songbird Sally-Anne test. (**A**) Student bird 1 learns how a teacher bird sings. Student bird 2 learns how the teacher and student 1 sing, respectively. (**B**) Student 2 notices that the teacher changes its song but student 1 does not know this, because it is absent (the Sally-Anne test). Student 2 can infer how student 1 behaves by distinguishing the difference between the internal states of itself and student 1. In these illustrations, a bird with a single internal model would be associated with a small child or autistic individual, while a bird with multiple internal models would be correspond to a healthy adult.

In summary, we have introduced a novel learning scheme that integrates Bayesian filtering and model selection to learn and deploy multiple generative models. We assumed that a switching variable selects a particular model to generate current sensory input (like switching to a particular radio channel from a repertoire of radio programs), while many alternative generative models are running in the background. To deal with the problem of context sensitive learning, the proposed scheme calculates the model plausibility (i.e. model evidence) of each generative model based on conditional free actions and updates parameters only in models with high evidence. Our synthetic agents were able to both learn and recognise different artificial and natural birdsongs. These results highlight the potential utility of equipping agents with multiple generative models to make inferences in context sensitive environments.

## METHODS

### Variational update scheme

The proposed variational update scheme is described in the Results. The further details are provided as Supplementary Methods S1 and S2.

### Songbird model

A generative model for bird song generation is defined as a two-layer hierarchical generative model, in which each layer has three hidden states 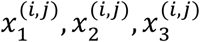 that generate a Lorenz attractor [32,33]. For simulations, we used several different teacher songs and the same number of internal models.

***For Fig. 3*** Two teacher songs were generated from two generative models with different parameter. A student was supposed to have two internal models. We defined a generative model according to [32,33]. Layer 1 has one hidden cause 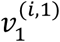 and one parameter 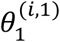, while layer 2 has no hidden cause nor parameter. Functions were defined as follows:

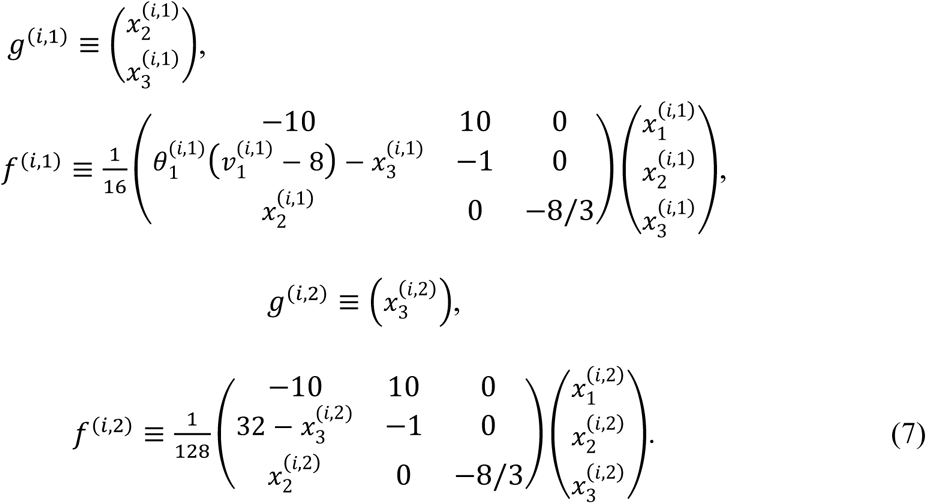

Different teacher songs used different parameter 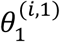. This parameter was learnt by a student without supervision. Initial states of latent variables were defined by *x*^(*i*,1)^ = (1,1,30)^0^ and *x*^(*i*,2)^ = (1,1,1)^*T*^, while the posterior expectations were initialised as **x**^(*i*,1)^ = (1,1,1)^*T*^ and **x**^(*i*,2)^ = (1,1,1)^*T*^. Training was repeated for 32 sessions. MATLAB source codes for this simulation are appended as Supplementary Source Codes. To run the simulation, one needs to install the open access academic software SPM (http://www.fil.ion.ucl.ac.uk/spm/software/).

***For Fig. 4*** We used six natural zebra finch songs as teacher songs and supposed a student that has six internal models. These models share the same structure while using different latent variables and parameters. We slightly changed the above generative model to deal with real zebra finch songs. Layer 1 has three hidden causes 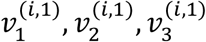 and 420-dimensional parameters (a 2 × 210 matrix *θ*^(*i*,1)^), while layer 2 has no hidden cause nor parameter. Functions were defined as follows:

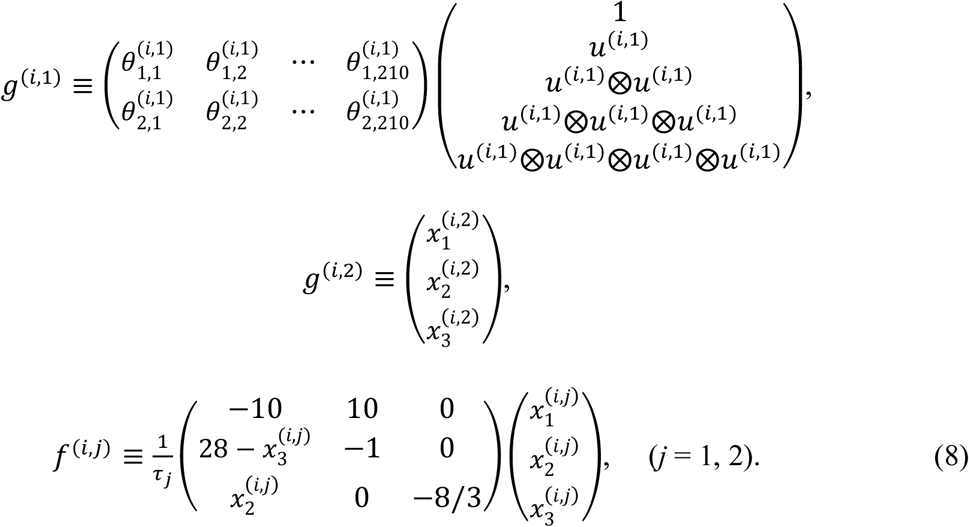

Note that 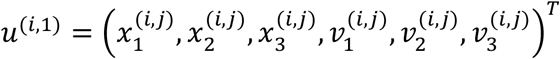 is a vector of latent variables and ⊗ expresses outer product (arranging all pair-wise products of elements of two vectors in vertical line except duplicate terms). We supposed that hidden causes *ν*^(*i*,1)^ has converged to *x*^(*i*,2)^ to simplify the simulation. The definition of *g*^(*i*,1)^ is to ensure it can express a general quartic function by a linear product of a 2 × 210 matrix and a 210-dimensional vector. These parameters were learnt by a student without supervision. When updating the posterior belief of hidden states, to avoid a divergence of variables induced by a large perturbation, we smoothed the posterior trajectory by adding small amount of components of the prior (i.e., a trajectory of a Lorenz attractor without perturbation).

The following procedure was applied before training: (1) initial states of **x**^(*i*,2)^, **x**^(*i*,2)^ and time constants *τ*_2_, *τ*_1_ were optimised to one of six teacher songs; (2) the posterior expectation of parameters **θ**^(*i*,1)^ were generated randomly; and (3) **θ**^(*i*,1)^ were modified by pretraining, in which each model randomly received one of six teacher songs, made inference and updated parameters without model selection for 18 sessions. Then, we tested response songs of a student to different teacher songs (Supplementary Movie S1). For training, we randomly selected one of six teacher songs and provided it to a student (Supplementary Movie S2). Training was repeated for 60 sessions. After training, we tested response songs again (Fig. 4 and Supplementary Movie S3).

Birdsong data used for Fig. 4 and Supplementary Movies were downloaded from http://ofer.sci.ccny.cuny.edu/song_database/zebra-finch-song-library-2015/view

This dataset was recorded by Tchernichovski group (see e.g., [42]). We treated the data accordingly. First, we got a spectrogram of song by Fourier transform with a 23.2-ms time window. As an analogy to a physiological model of vocal coda that generates a birdsong by sequences of power and tone (frequency) of voice [47], we defined the leading frequency (*s*_2_) and the amplitude (*s*_1_) of a song by the mode of frequency and its power for each time step, respectively. They were normalised and introduced as sensory inputs *s* = (*s*_1_, *s*_2_)*^T^*.

## Acknowledgements

This work was supported by RIKEN Brain Science Institute (T.I.) and Tateisi Science and Technology Foundation (T.I.). T.P. is supported by the Rosetrees Trust (Award Number 173346). K.J.F. is funded by a Wellcome Trust Principal Research Fellowship (Ref: 088130/Z/09/Z). The funders had no role in study design, data collection and analysis, decision to publish, or preparation of the manuscript.

## Author Contributions

Conceptualized the free-energy principle: K.J.F. Conceived and designed the method using the multiple internal models: T.I. Performed the simulations: T.I. Wrote the paper: T.I., T.P., and K.J.F.

