## Supplementary Materials for "Social intelligence model with multiple internal models"

#### Supplementary Tables

Table S1: Glossary of expressions.

| **Expression** | **Description** |
|---:|---|
| $\tilde{s} \in \mathbb{R}^{\bullet}$ | Sensory inputs (teacher song) |
| $m_i$ | Model $i$ (a specific model structure) |
| $m_c$ | Currently selected model |
| $\mathcal{M}$ | Set of model indices |
| $\tilde{x}^{(i,j)} \in \mathbb{R}^{\bullet}$ | Hidden states of the $j$-th layer in model $i$ |
| $\tilde{v}^{(i,j)} \in \mathbb{R}^{\bullet}$ | Hidden causes of the $j$-th layer in model $i$ |
| $u^{(i)} \equiv (\tilde{x}^{(i,1)}, \tilde{v}^{(i,1)}, \tilde{x}^{(i,2)}, \tilde{v}^{(i,2)}) \in \mathbb{R}^{\bullet}$ | Latent variables of model $i$ |
| $\theta^{(i)} \equiv (\theta^{(i,1)}, \theta^{(i,2)}) \in \mathbb{R}^{\bullet}$ | Parameters of model $i$ |
| $P(i = c) \equiv \gamma_i \in \{0, 1\}$ with $\sum_{i \in \mathcal{M}} \gamma_i = 1$ | Sufficient statistics of switcher state |
| $P(\gamma) \equiv Cat(\Gamma)$ | Prior distribution of $\gamma = (\gamma_1, \gamma_2, \dots)$ |
| $\mathbf{u}^{(i)} \in \mathbb{R}^{\bullet}$ | Posterior expectation of latent variables (mean of $q(u^{(i)})$) |
| $\boldsymbol{\theta}^{(i)} \in \mathbb{R}^{\bullet}$ | Posterior expectation of parameters (mean of $q(\theta^{(i)})$) |
| $Q(i = c) \equiv \boldsymbol{\gamma}_i \in [0, 1]$ with $\sum_{i \in \mathcal{M}} \boldsymbol{\gamma}_i = 1$ $\iff Q(\gamma) \equiv Cat(\boldsymbol{\gamma})$ | Posterior expectation of switcher state (model plausibility) |
| $\tilde{\omega}^{(i,j)}, \tilde{z}^{(i,j)} \in \mathbb{R}^{\bullet}$ | Background noises |
| $\tilde{g}^{(i,j)}, \tilde{f}^{(i,j)} \in \mathbb{R}^{\bullet}$ | Functions |
| $a \in \mathbb{R}^{\bullet}$ | Action (student song) |
| $\mathrm{E}_{p(x)}[\bullet(x)] \equiv \int \bullet(x) p(x) dx$ | Expectation of $\bullet(x)$ over $p(x)$ |
| $\mathcal{D}_{KL}[q(\bullet) \Vert p(\bullet)] \equiv \mathrm{E}_{q(\bullet)}\left[\log \frac{q(\bullet)}{p(\bullet)}\right]$ | Kullback-Leibler divergence between $q(\bullet)$ and $p(\bullet)$ |
| $U_i(\tilde{s}, u^{(i)}, \theta^{(i)}) \equiv -\log p(\tilde{s}, u^{(i)} \vert \theta^{(i)}, m_i)$ | Internal energy under model $i$ |
| $F_i(t) \equiv U_i(\tilde{s}, u^{(i)}, \theta^{(i)}) + \mathrm{E}_{q(u^{(i)})}[\log q(u^{(i)})]$ | Conditional free energy under model $i$ |
| $F(t) \equiv \mathrm{E}_{Q(i=c)}[F_i(t)] \equiv \sum_{i \in \mathcal{M}} \boldsymbol{\gamma}_i F_i(t)$ | Total free energy |
| $\overline{F}_i \equiv \int_0^T F_i(t) dt$ | Conditional free action under model $i$ |
| $\overline{F} \equiv \mathrm{E}_{Q(i=c)}\left[\overline{F}_i\right] + \mathcal{D}_{KL}\left[Q(\gamma) \Vert P(\gamma)\right]$ $+ \sum_{i \in \mathcal{M}} \mathcal{D}_{KL}\left[q(\theta^{(i)}) \Vert p(\theta^{(i)} \vert m_i)\right]$ | Total free action |

### Supplementary Figures

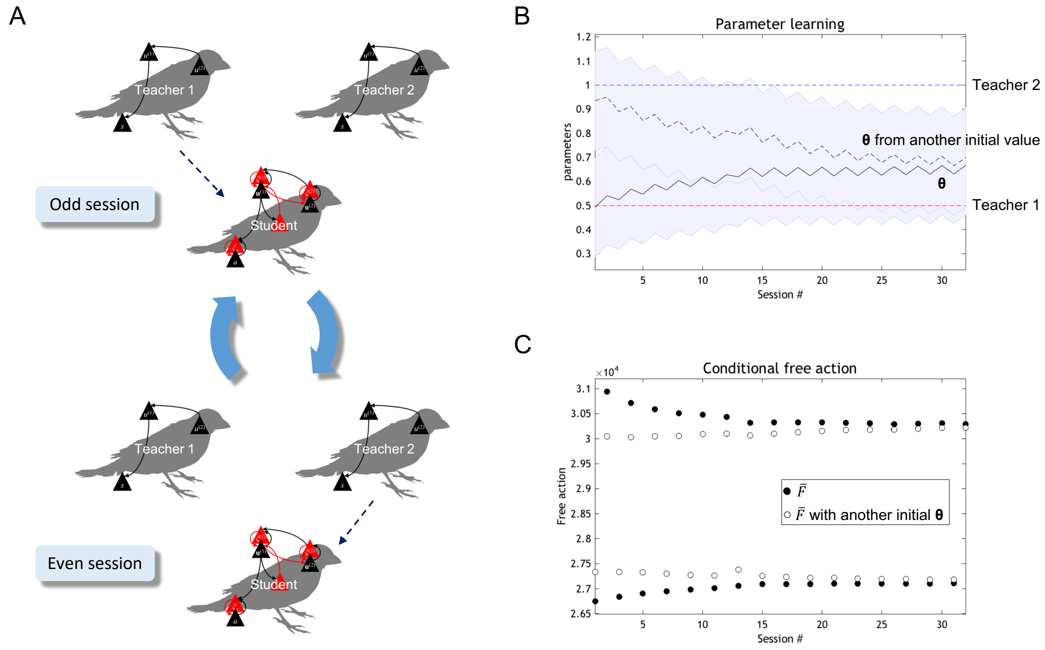


**Figure S1.** A schematic illustrating an experimental procedure (A) and simulation results of a synthetic bird with a single generative model (B,C). (**A**) Experimental procedure. Teacher bird 1 sings in odd sessions while teacher bird 2 sings in even sessions. Our synthetic bird (student) listens either song in turn. (**B**) Trajectory of the posterior expectation of a parameter ($\boldsymbol{\theta}$) of the student that employs a single generative model (black sold curve). A black dashed curve shows another trajectory where $\boldsymbol{\theta}$ started from a different initial value. Shaded areas indicate the standard deviation. Red and blue dashed lines express the true parameter of teacher 1 and 2, respectively. The student tried to infer either parameter of teacher 1 or 2, but it failed to learn either parameter even its posterior belief was initialized to the same value as either teacher 1 or 2's parameter. This is because the student inferred the intermediate value of teacher 1 and 2's parameters. (**C**) Transition of free action (filled circles). Open circles show transition of free energy with another initial $\boldsymbol{\theta}$.

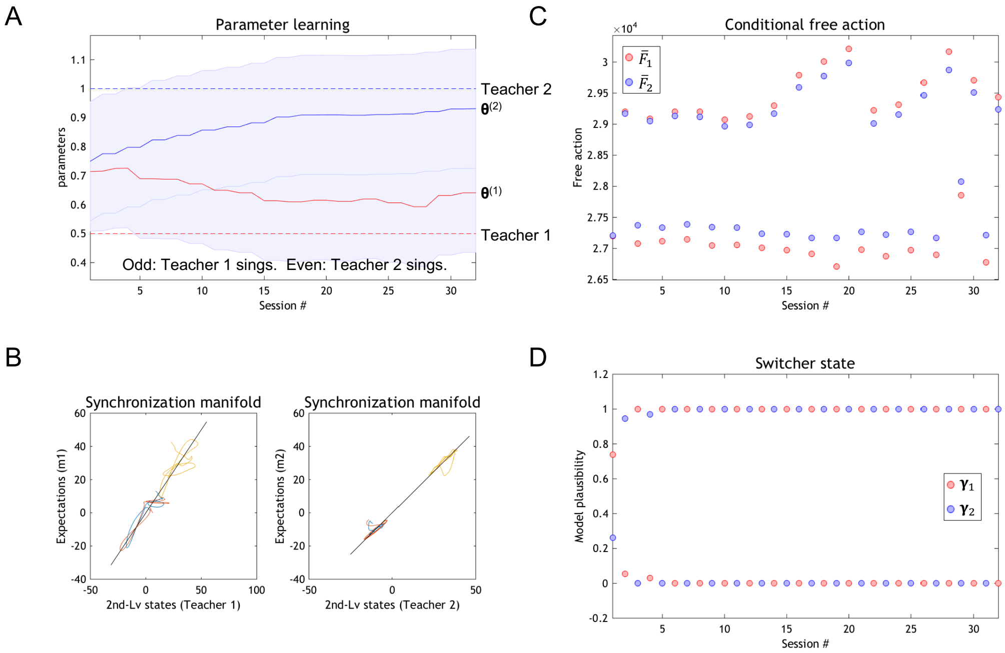


**Figure S2.** Simulation results when learning two birdsongs using multiple generative models. Simulation setup and layout are the same as Fig. 3, but initial hidden states of teacher 1 and 2 were not reset for each session. This yielded the chaotic dynamics in their songs. Even in this case, a student bird employing the proposed scheme could learn from two distinct teachers.

#### Supplementary Methods

##### S1 Generative model

Formally, the multiple generative model is defined as the following. Hierarchical Bayesian filtering supposes a model that consists of latent variables $u$ (a set of hidden states $x$ and hidden causes $v$) and parameters $\theta$, and infers their approximate probability (recognition) densities. To extend this for a multiple-model version, we express the $i$-th generative model consisting of two layers as $m_i$ with $i \in \mathcal{M} \equiv \{1, 2, 3, \dots\}$. This $m_i$ indicates a specific model structure including certain forms of functions and dimensions of latent variables and parameters. Let $\tilde{s}$ be sensory inputs (i.e., teacher song) generated by $m_i$, and $\tilde{x}^{(i,j)}$, $\tilde{v}^{(i,j)}$, and $\theta^{(i,j)}$ be hidden states, hidden causes, and parameters in the $j$-th layer ($j = 1, 2$) of $m_i$, respectively. The tilde over a symbol denotes a set of a variable and its time derivatives $\tilde{s} \equiv (s, s', s'', \dots)$. Throughout this paper, $i$ indices the model while $j$ indices the level of layers. The $i$-th generative model is given by Eq. (1) in the main text. The corresponding probabilities are defined as Gaussian distributions and written as

$$
\begin{aligned}
\tilde{s} &= \tilde{g}^{(i,1)} + \tilde{\omega}^{(i,1)} &&\Longleftrightarrow\quad p\left(\tilde{s} \;\middle|\; \tilde{x}^{(i,1)}, \tilde{v}^{(i,1)}, \theta^{(i,1)}, m_i\right) \equiv \mathcal{N}\left[\tilde{s}; \tilde{g}^{(i,1)}, \Pi_v^{(i,1)}\right],\\
D\tilde{x}^{(i,1)} &= \tilde{f}^{(i,1)} + \tilde{z}^{(i,1)} &&\Longleftrightarrow\quad p\left(\tilde{x}^{(i,1)} \;\middle|\; \tilde{v}^{(i,1)}, \theta^{(i,1)}, m_i\right) \equiv \mathcal{N}\left[D\tilde{x}^{(i,1)}; \tilde{f}^{(i,1)}, \Pi_x^{(i,1)}\right],\\
\tilde{v}^{(i,1)} &= \tilde{g}^{(i,2)} + \tilde{\omega}^{(i,2)} &&\Longleftrightarrow\quad p\left(\tilde{v}^{(i,1)} \;\middle|\; \tilde{x}^{(i,2)}, \tilde{v}^{(i,2)}, \theta^{(i,2)}, m_i\right) \equiv \mathcal{N}\left[\tilde{v}^{(i,1)}; \tilde{g}^{(i,2)}, \Pi_v^{(i,2)}\right],\\
D\tilde{x}^{(i,2)} &= \tilde{f}^{(i,2)} + \tilde{z}^{(i,2)} &&\Longleftrightarrow\quad p\left(\tilde{x}^{(i,2)} \;\middle|\; \tilde{v}^{(i,2)}, \theta^{(i,2)}, m_i\right) \equiv \mathcal{N}\left[D\tilde{x}^{(i,2)}; \tilde{f}^{(i,2)}, \Pi_x^{(i,2)}\right],\\
& && \qquad p\left(\tilde{v}^{(i,2)} \;\middle|\; m_i\right) \equiv \mathcal{N}\left[\tilde{v}^{(i,2)}; \tilde{\eta}^{(i)}, \Pi_\eta^{(i)}\right],\\
& && \qquad p\left(\theta^{(i,j)} \;\middle|\; m_i\right) \equiv \mathcal{N}\left[\theta^{(i,j)}; \Theta^{(i,j)}, \Pi_\theta^{(i,j)}\right], \quad (j = 1, 2).
\end{aligned}
\tag{S1}
$$

Note that $\tilde{\omega}^{(i,j)} \sim \mathcal{N}\left[\omega^{(i,j)}; 0, \Pi_v^{(i,j)}\right]$ and $\tilde{z}^{(i,j)} \sim \mathcal{N}\left[z^{(i,j)}; 0, \Pi_x^{(i,j)}\right]$ are background Gaussian noises, $\tilde{g}^{(i,j)} \equiv \tilde{g}^{(i,j)}\left(\tilde{x}^{(i,j)}, \tilde{v}^{(i,j)}, \theta^{(i,j)}\right)$ and $\tilde{f}^{(i,j)} \equiv \tilde{f}^{(i,j)}\left(\tilde{x}^{(i,j)}, \tilde{v}^{(i,j)}, \theta^{(i,j)}\right)$ are arbitrary functions of $\tilde{x}^{(i,j)}$ and $\tilde{v}^{(i,j)}$ parameterized by $\theta^{(i,j)}$, $\Pi_v^{(i,j)}$ and $\Pi_x^{(i,j)}$ are precision matrices, and $p(\tilde{v}^{(i,2)}|m_i)$ and $p(\theta^{(i,j)}|m_i)$ are Gaussian priors parameterized by the mean and the precision matrix. For simplification of notation, we suppose $u^{(i,j)} \equiv (\tilde{x}^{(i,j)}, \tilde{v}^{(i,j)})$ as latent variables, $u^{(i)} \equiv (u^{(i,1)}, u^{(i,2)})$ as a set of latent variables in all layers of $m_i$, and $\theta^{(i)} \equiv (\theta^{(i,1)}, \theta^{(i,2)})$ as a set of parameters in all layers of $m_i$. By multiplying all equations in the right side of Eq. (S1), the $i$-th generative model is expressed as

$$
\begin{aligned}
p\left(\tilde{s}, u^{(i)}, \theta^{(i)} \;\middle|\; m_i\right) = p\left(\tilde{s}^{(i)} \;\middle|\; \tilde{x}^{(i,1)}, \tilde{v}^{(i,1)}, \theta^{(i,1)}, m_i\right) p\left(\tilde{v}^{(i,1)} \;\middle|\; \tilde{x}^{(i,2)}, \tilde{v}^{(i,2)}, \theta^{(i,2)}, m_i\right) p\left(\tilde{v}^{(i,2)} \;\middle|\; m_i\right)\\
\cdot \prod_{j=1,2} p\left(\tilde{x}^{(i,j)} \;\middle|\; \tilde{v}^{(i,j)}, \theta^{(i,j)}, m_i\right) p\left(\theta^{(i,j)} \;\middle|\; m_i\right)
\end{aligned}
\tag{S2}
$$

as shown on the top in Fig. 1.

**Sensory inputs** The sensory input that an agent (i.e., a student bird) actually receives is selected by one of models in $\mathcal{M} \equiv \{1, 2, \ldots\}$, where we index the currently selected model by $c$. The sensory input is expressed by the sum of the product of the conditional probability of $\tilde{s}$ under each model and the probability of each model being selected,

$$p(\tilde{s} \mid m_c) \equiv \mathrm{E}_{P(i=c)}\left[p(\tilde{s} \mid m_i)\right] \equiv \sum_{i \in \mathcal{M}} P(i = c) p(\tilde{s} \mid m_i) \equiv \sum_{i \in \mathcal{M}} \gamma_i p(\tilde{s} \mid m_i) \equiv \prod_{i \in \mathcal{M}} p(\tilde{s} \mid m_i)^{\gamma_i}. \tag{S3}$$

Note that the sufficient statistics $\gamma_i \equiv P(i = c) \in \{0, 1\}$ is a binary value that satisfies $\sum_{i \in \mathcal{M}} \gamma_i = 1$ by design, where only $\gamma_c$ takes one while others being zero. In this design, $\gamma_i$ plays a role of a switcher that switches which model has generated the current sensory input, while all models are running in the background. The probability of $\gamma = (\gamma_1, \gamma_2, \ldots)$ follows a categorical prior distribution $P(\gamma) = Cat(\Gamma)$.

Interestingly, this definition of multiple models and a switcher is slightly different from supposing one big generative model. This idea is rather an assumption that an agent has a set of hypotheses (generative models) about sensory inputs to select an appropriate one for different contexts. Different generative models are running independently from each other. They are interacted only via sensory inputs through a model switching scheme, but this interaction does not change their latent variables or parameters. Since each model can be considered independently, each model can have different structures and dimensions, while in this study we assume models based on the same model structure and dimension with different latent variables and parameters.

**Free energy and free action** The negative log of $p(\tilde{s} \mid m_c)$ gives surprise of sensory inputs and free energy is defined as an upper bound of surprise [14,15,30]. In this study, we slightly modify the derivation of free energy to include a model selection mechanism. First, we show that Bayesian model averaging of the conditional surprises provides an upper bound of surprise. Suppose $Q(i = c) \equiv \boldsymbol{\gamma}_i \in [0, 1]$ with $\sum_{i \in \mathcal{M}} \boldsymbol{\gamma}_i = 1$ is the posterior expectation of the switcher state. This is equivalent to a categorical posterior distribution of $\gamma$ given by $Q(\gamma) = Cat(\boldsymbol{\gamma})$. Since model $c$ is selected, the following inequality holds from the non-negativity of Kullback-Leibler divergence [38],

$$\begin{aligned}\mathrm{E}_{p(\tilde{s} \mid m_c)}\left[\log p(\tilde{s} \mid m_c) - \sum_{i \in \mathcal{M}} \boldsymbol{\gamma}_i \log p(\tilde{s} \mid m_i)\right] &= \sum_{i \in \mathcal{M}} \boldsymbol{\gamma}_i \mathrm{E}_{p(\tilde{s} \mid m_c)}\left[\log p(\tilde{s} \mid m_c) - \log p(\tilde{s} \mid m_i)\right] \\ &= \sum_{i \in \mathcal{M}} \boldsymbol{\gamma}_i \mathcal{D}_{KL}\left[p(\tilde{s} \mid m_c) \,||\, p(\tilde{s} \mid m_i)\right] \geq 0,\end{aligned} \tag{S4}$$

where $-\log p(\tilde{s} \mid m_i)$ is a conditional surprise under model $i$. The expectation over sensory inputs $\mathrm{E}_{p(\tilde{s})}[\bullet]$ can be approximately calculated using time average of surprise under the ergodic assumption. From Eq. (S4), we have

$$\mathrm{E}_{p(\tilde{s} \mid m_c)Q(i=c)}\left[-\log p(\tilde{s} \mid m_i)\right] = \mathrm{E}_{p(\tilde{s} \mid m_c)}\left[-\sum_{i \in \mathcal{M}} \boldsymbol{\gamma}_i \log p(\tilde{s} \mid m_i)\right] \geq \mathrm{E}_{p(\tilde{s} \mid m_c)}\left[-\log p(\tilde{s} \mid m_c)\right], \tag{S5}$$

and also approximately have

$$\mathrm{E}_{Q(i=c)}\left[\int_0^T -\log p(\tilde{s}_t \mid m_i)dt\right] = \sum_{i\in\mathcal{M}} \boldsymbol{\gamma}_i \int_0^T -\log p(\tilde{s}_t \mid m_i)dt \geq \int_0^T -\log p(\tilde{s}_t \mid m_c)dt, \tag{S6}$$

where $\tilde{s}_t$ is a sampling of $\tilde{s}$ at time $t$ and $T$ is the measurement time. Finally, we define total free action (the path integral of free energy) as an upper bound of $\mathrm{E}_{Q(i=c)}\left[\int_0^T -\log p(\tilde{s}_t \mid m_i)dt\right]$,

$$\begin{aligned}\overline{F} &\equiv \mathrm{E}_{Q(i=c)}\left[\int_0^T \left(-\log p(\tilde{s}_t \mid m_i) + \mathrm{E}_{q(\theta^{(i)})}\left[\mathcal{D}_{KL}\left[q(u_t^{(i)})\|p(u_t^{(i)} \mid \tilde{s}_t, \theta^{(i)}, m_i)\right]\right]\right)dt\right] \\ &\qquad + \mathcal{D}_{KL}\left[Q(\gamma)\|P(\gamma)\right] + \sum_{i\in\mathcal{M}} \mathcal{D}_{KL}\left[q(\theta^{(i)})\|p(\theta^{(i)} \mid m_i)\right] \\ &= \mathrm{E}_{Q(i=c)}\left[\overline{F}_i + \log\boldsymbol{\gamma}_i - \log\Gamma_i\right] + \sum_{i\in\mathcal{M}} \mathrm{E}_{q(\theta^{(i)})}\left[\log q(\theta^{(i)}) - \log p(\theta^{(i)} \mid m_i)\right] \\ &\geq \mathrm{E}_{Q(i=c)}\left[\int_0^T -\log p(\tilde{s}_t \mid m_i)dt\right].\end{aligned} \tag{S7}$$

Note that $q(u_t^{(i)})$ and $q(\theta^{(i)})$ are the posterior densities of latent variables and parameters under model $i$, respectively. In this expression, the total free energy is defined as the weighed sum of conditional free actions $\overline{F}_1, \overline{F}_2, \ldots$ plus Kullback-Leibler divergence (i.e., complexity) of between the prior and posterior of the switcher state and parameters. Each conditional free action is defined by

$$\overline{F}_i \equiv \int_0^T \mathrm{E}_{q(u_t^{(i)})q(\theta^{(i)})}\left[-\log p(\tilde{s}_t, u_t^{(i)} \mid \theta^{(i)}, m_i) + \log q(u_t^{(i)})\right]dt \equiv \int_0^T F_i(t)dt, \tag{S8}$$

where $F_i(t)$ is a conditional free energy and expressed in the following three ways,

$$\begin{aligned}F_i(t) &\equiv -\log p\left(\tilde{s} \mid m_i\right) + \mathrm{E}_{q(\theta^{(i)})}\left[\mathcal{D}_{KL}\left[q(u^{(i)}) \parallel p(u^{(i)} \mid \tilde{s}, \theta^{(i)}, m_i)\right]\right] \\ &= \mathrm{E}_{q(u^{(i)})q(\theta^{(i)})}\left[-\log p\left(\tilde{s} \mid u^{(i)}, \theta^{(i)}, m_i\right)\right] + \mathrm{E}_{q(\theta^{(i)})}\left[\mathcal{D}_{KL}\left[q(u^{(i)}) \parallel p(u^{(i)} \mid \theta^{(i)}, m_i)\right]\right] \\ &= \mathrm{E}_{q(u^{(i)})q(\theta^{(i)})}\left[-\log p\left(\tilde{s}, u^{(i)} \mid \theta^{(i)}, m_i\right) + \log q(u^{(i)})\right].\end{aligned} \tag{S9}$$

In the last line, the first term is the negative log of Eq. (S2) divided by $p(\theta^{(i)} \mid m_i)$ and will be referred to as internal energy under model $i$, $U_i(\tilde{s}, u^{(i)}, \theta^{(i)}) \equiv -\log p\left(\tilde{s}, u^{(i)} \mid \theta^{(i)}, m_i\right)$. Note that $F(t) \equiv \mathrm{E}_{Q(i=c)}[F_i(t)] \equiv \sum_{i\in\mathcal{M}} \boldsymbol{\gamma}_i F_i(t)$ denotes total free energy.

**Posteriors** From Laplace assumption, the posterior density of latent variables $u^{(i)}$ is approximated as a Gaussian distribution $q(u^{(i)}) = \mathcal{N}[u^{(i)}; \mathbf{u}^{(i)}, P_u^{(i)}]$ with an expectation (or mode) vector $\mathbf{u}^{(i)}$ and a precision matrix $P_u^{(i)}$. The posterior density of parameters $\theta^{(i)}$ is approximated as a Gaussian distribution $q(\theta^{(i)}) = \mathcal{N}[\theta^{(i)}; \boldsymbol{\theta}^{(i)}, P_\theta^{(i)}]$ with an expectation vector $\boldsymbol{\theta}^{(i)}$ and a precision matrix $P_\theta^{(i)}$. As described above, the posterior distribution of the switcher state (i.e., model plausibility) has been defined as a categorical distribution $Q(i = c) \equiv \boldsymbol{\gamma}_i$ with $\sum_{i \in \mathcal{M}} \boldsymbol{\gamma}_i = 1$, which is equivalent to $Q(\gamma) = Cat(\boldsymbol{\gamma})$.

###### S2 Update rules

Updates of posteriors of latent variables, the switcher state, and parameters are conducted in inference, model selection, and learning steps, respectively. In simulation, these three steps are repeated in order for each session, but one can also consider that they are updated simultaneously with different time scales. In what follows, we formally derive update rules from the minimization of free energy or free action.

**Inference** The optimal $q(u^{(i)})$ is obtained by solving the variation of $F_i$, $\delta F_i / \delta q(u^{(i)}) = \mathrm{E}_{q(\theta^{(i)})}\left[U_i(\tilde{s}, u^{(i)}, \theta^{(i)})\right] + \log q(u^{(i)}) + \text{const.} = 0$. To satisfy this condition, the density should be $q(u^{(i)}) \propto \exp\left[-\mathrm{E}_{q(\theta^{(i)})}\left[U_i(\tilde{s}, u^{(i)}, \theta^{(i)})\right]\right] \approx \exp[-U_i(\tilde{s}, u^{(i)}, \boldsymbol{\theta}^{(i)})]$, where $U_i(\tilde{s}, u^{(i)}, \boldsymbol{\theta}^{(i)})$ is the zero-th order approximation of the variational energy for latent variables. The dynamics of $\mathbf{u}^{(i)}$ are given by $\dot{\mathbf{u}}^{(i)} = D\mathbf{u}^{(i)}$ with a derivative operator $D$. When $\mathbf{u}^{(i)}$ has been optimized, the path of the mode $\dot{\mathbf{u}}^{(i)}$ should be equal to the mode of the path $D\mathbf{u}^{(i)}$, $\dot{\mathbf{u}}^{(i)} = D\mathbf{u}^{(i)}$ in addition to minimize $U_i$ (see [14,28] for details). Thus, the gradient descent rule for $\mathbf{u}^{(i)}$ that minimizes $F_i$ is given by

$$\dot{\mathbf{u}}^{(i)} - D\mathbf{u}^{(i)} \propto -\frac{\partial}{\partial u^{(i)}} U_i(\tilde{s}, u^{(i)}, \boldsymbol{\theta}^{(i)})\Big|_{u^{(i)}=\mathbf{u}^{(i)}} = -\frac{\partial}{\partial \mathbf{u}^{(i)}} U_i(\tilde{s}, \mathbf{u}^{(i)}, \boldsymbol{\theta}^{(i)}) \approx -\frac{\partial}{\partial \mathbf{u}^{(i)}} F_i(t) \tag{S10}$$

and $P_u^{(i)}$ that minimizes $F_i$ is given by

$$P_u^{(i)} = \frac{\partial^2}{\partial (u^{(i)})^2} U_i(\tilde{s}, u^{(i)}, \boldsymbol{\theta}^{(i)})\Big|_{u^{(i)}=\mathbf{u}^{(i)}} \approx \frac{\partial^2}{\partial (\mathbf{u}^{(i)})^2} F_i(t) \tag{S11}$$

Eq. (S10) is the same as Eq. (3) in the main text.

**Model selection** This step performs online Bayesian model selection analogous to post hoc Bayesian model selection [39]. Since the posterior expectations of the switcher state are discrete variables, this step is similar to Markov decision process model [18,40]. From Eq. (S7), a condition to be an extreme value is solved as $\delta \overline{F} / \delta \boldsymbol{\gamma}_i = \overline{F}_i + \log \boldsymbol{\gamma}_i - \log \Gamma_i + \text{const.} = 0$. Therefore, we obtain $\boldsymbol{\gamma}_i$ that minimizes $\overline{F}$ as

$$\boldsymbol{\gamma}_i = \sigma\left(-\overline{F}_i\left(\boldsymbol{\theta}^{(i)}\right) + \log \Gamma_i\right), \tag{S12}$$

where $\sigma(\bullet_i)$ is a softmax function defined by $\sigma(\bullet_i) \equiv \exp(\bullet_i) / \sum_{k \in \mathcal{M}} \exp(\bullet_k)$. This $\boldsymbol{\gamma}_i$ expresses the model plausibility of model $i$. When we suppose the prior distribution $P(\gamma)$ as flat (i.e., a uniform distribution), Eq. (S12) becomes Eq. (5) in the main text.

**Learning** Estimation of parameters is based on conventional gradient descent approach. To satisfy $\delta \overline{F} / \delta q(\theta^{(i)}) = \boldsymbol{\gamma}_i \int_0^T \mathrm{E}_{q(\tilde{u}_t^{(i)})}[U_i(\tilde{s}_t, \tilde{u}_t^{(i)}, \theta^{(i)})]dt +$

$\log q(\theta^{(i)}) - \log p(\theta^{(i)}|m_i) + \text{const.} = 0$, the density should be $q(\theta^{(i)}) \propto \exp\left[-\boldsymbol{\gamma}_i \int_0^T \mathrm{E}_{q(u_t^{(i)})}\left[U_i(\tilde{s}_t, \tilde{u}_t^{(i)}, \theta^{(i)})\right] dt + \log p(\theta^{(i)}|m_i)\right] \approx \exp\left[-\boldsymbol{\gamma}_i \int_0^T U_i(\tilde{s}_t, \tilde{\mathbf{u}}_t^{(i)}, \theta^{(i)}) dt + \log p(\theta^{(i)}|m_i)\right]$, where $\int_0^T U_i(\tilde{s}_t, \tilde{\mathbf{u}}_t^{(i)}, \theta^{(i)}) dt$ is the approximate variational action for parameters. The gradient descent (steepest descent) rule for $\boldsymbol{\theta}^{(i)}$ that minimizes $\overline{F}$ is given by

$$\begin{aligned}\dot{\boldsymbol{\theta}}^{(i)} &\propto -\frac{\partial}{\partial \theta^{(i)}}\left(\boldsymbol{\gamma}_i \int_0^T U_i(\tilde{s}_t, \tilde{\mathbf{u}}_t^{(i)}, \theta^{(i)}) dt - \log p(\theta^{(i)}|m_i)\right)\Bigg|_{\theta^{(i)}=\boldsymbol{\theta}^{(i)}} \\ &= -\frac{\partial}{\partial \boldsymbol{\theta}^{(i)}}\left(\boldsymbol{\gamma}_i \int_0^T U_i(\tilde{s}_t, \tilde{\mathbf{u}}_t^{(i)}, \boldsymbol{\theta}^{(i)}) dt - \log p(\boldsymbol{\theta}^{(i)}|m_i)\right) \approx -\frac{\partial}{\partial \boldsymbol{\theta}^{(i)}}\left(\boldsymbol{\gamma}_i \overline{F}_i\left(\boldsymbol{\theta}^{(i)}\right) - \log p\left(\boldsymbol{\theta}^{(i)}|m_i\right)\right)\end{aligned} \tag{S13}$$

and $P_\theta^{(i)}$ that minimizes $\overline{F}$ is given by

$$\begin{aligned}\dot{P}_\theta^{(i)} &\propto -P_\theta^{(i)} + \frac{\partial^2}{\partial (\theta^{(i)})^2}\left(\boldsymbol{\gamma}_i \int_0^T U_i(\tilde{s}_t, \tilde{\mathbf{u}}_t^{(i)}, \theta^{(i)}) dt - \log p(\theta^{(i)}|m_i)\right)\Bigg|_{\theta^{(i)}=\boldsymbol{\theta}^{(i)}} \\ &\approx -P_\theta^{(i)} + \frac{\partial}{\partial (\boldsymbol{\theta}^{(i)})^2}\left(\boldsymbol{\gamma}_i \overline{F}_i\left(\boldsymbol{\theta}^{(i)}\right) - \log p\left(\boldsymbol{\theta}^{(i)}|m_i\right)\right)\end{aligned} \tag{S14}$$

When we suppose the prior distribution of parameters $P(\theta^{(i)}|m_i)$ as flat, Eq. (S13) becomes Eq. (6) in the main text.

Accordingly, we obtain posterior densities of latent variables and parameters, and posterior expectations of the switcher state that give the minimum of the total free action. The difference in learning rate mediated by the model plausibility enables that only parameters in the most plausible models are updated while parameters in the remaining models are maintained in a winner-takes-all manner. Whereas, latent variables in all models are updated with a fixed update rate. Therefore, inference occurs for all generative models, while learning occurs only for the most plausible generative models. This mechanism enables the agent to make inference and learning with several different generative models.

**Action** Action $a$ is generated to minimize the total free energy $F(t) \equiv \sum_{i \in \mathcal{M}} \boldsymbol{\gamma}_i F_i(t)$, which is given by $\dot{a} \propto -\partial F / \partial a$. Suppose the absence of sensory input from the environment, action directly induces sensory input $\tilde{s} = a$, and all internal models use the same precision matrix. In this special case, the optimal action is approximately solved as

$$\dot{a} \propto -\frac{\partial F}{\partial a} \approx \sum_{i \in \mathcal{M}} \boldsymbol{\gamma}_i g^{(i,1)}\left(\mathbf{u}^{(i,1)}, \boldsymbol{\theta}^{(i,1)}\right). \tag{S15}$$
